# Global Transcriptomics and Targeted Metabolite Analysis Reveal the Involvement of the AcrAB Efflux Pump in Physiological Functions by Exporting Signaling Molecules in *Photorhabdus laumondii*

**DOI:** 10.1101/2025.02.10.637478

**Authors:** Linda Hadchity, Anne Lanois-Nouri, Adrien Chouchou, David Roche, Jessica Houard, Noémie Claveyroles, Alexandra Dauvé, Jacques Imbert, Maxime Gualtieri, Alain Givaudan, Alyssa Carré-Mlouka, Ziad Abi Khattar

**Author notes:** Correspondence and reprints (A. Givaudan), (A. Carré-Mlouka) and (Z. Abi Khattar). (L. Hadchity), (A. Lanois-Nouri), (A. Chouchou), (D. Roche) (J. Houard), (N. Claveyroles); (A. Dauvé); (J. Imbert); (M. Gualtieri).

## Abstract

In Gram-negative bacteria, resistance-nodulation-division-type efflux pumps, particularly AcrAB-TolC, play a critical role in mediating resistance to antimicrobial agents and toxic metabolites, contributing to multidrug resistance. *Photorhabdus laumondii* is an entomopathogenic bacterium that has garnered significant interest due to its production of bioactive specialized metabolites with anti-inflammatory, antimicrobial, and scavenger deterrents properties. In previous work, we demonstrated that AcrAB confers self-resistance to stilbenes in *P. laumondii* TT01. Here, we explore the pleiotropic effects of AcrAB in this bacterium. RNA sequencing of Δ*acrA* compared to wild-type revealed growth-phase-specific gene regulation, with stationary-phase cultures showing significant downregulation of genes involved in stilbenes, fatty acid, and anthraquinone pigment biosynthesis, as well as genes related to cellular clumping and fimbrial pilin formation. Genes encoding putative LuxR regulators, type VI secretion systems, two-partner secretion systems, and contact dependent growth inhibition systems were upregulated in Δ*acrA*. Additionally, exponential-phase cultures revealed reduced expression of genes related to motility in Δ*acrA*. The observed transcriptional changes were consistent with phenotypic assays, demonstrating that the Δ*acrA* mutant had altered bioluminescence and defective orange pigmentation due to disrupted anthraquinone production. These findings confirm the role of stilbenes as signaling molecules involved in gene expression, thereby shaping these phenotypes. Furthermore, we showed that AcrAB contributes to swarming and swimming motilities independently of stilbenes. Collectively, these results highlight that disrupting *acrAB* causes transcriptional and metabolic dysregulation in *P. laumondii*, likely by impeding the export of key signaling molecules such as stilbenes, which may serve as a ligand for global transcriptional regulators.

**Importance:** Recent discoveries have highlighted *Photorhabdus laumondii* as a promising source of novel anti-infective compounds, including non-ribosomal peptides and polyketides. One key player in the self-resistance of this bacterium to stilbene derivatives is the AcrAB-TolC complex, which is also a well-known contributor to multidrug-resistance. Here, we demonstrate the pleiotropic effects of the AcrAB efflux pump in *P. laumondii* TT01, impacting secondary metabolite biosynthesis, motility, and bioluminescence. These effects are evident at transcriptional, metabolic, and phenotypic levels and are likely mediated by the efflux of signaling molecules such as stilbenes. These findings shed light on the multifaceted roles of efflux pumps and open avenues to better explore the complexity of RND pump-mediated signaling pathways in bacteria, thereby aiding combat multidrug resistant infections.

## Introduction

The Gram-negative bacterium *Photorhabdus laumondii* TT01 is a bioluminescent and motile entomopathogen that forms a mutualistic relationship with soil-dwelling nematodes of the *Heterorhabditidae* family [1, 2]. This bacterium-nematode complex is highly effective at infecting and killing insects by septicemia and toxemia, with death occurring within 24 to 48 h post-infection [1, 3, 4]. Recently, entomopathogenic bacteria such as *Photorhabdus* have garnered considerable interest in both fundamental and applied research, owing to the extensive array of bioactive specialized metabolites they produce during the saprophytic phase within insect cadavers [5–8]. The study of these compounds provide valuable insights into medicine, microbial ecology, and biochemistry [9, 10]. Many of them, including stilbene derivatives (STs), carbapenem-like compounds, odilorhabdins, indole, and anthraquinones (AQs), function to inhibit bacterial and fungal contamination and/or deter scavengers in insect cadavers [6, 8, 11–14].

In *Photorhabdus*, secondary metabolite production is regulated by several common transcriptional regulators, including HexA, an ortholog of LrhA in *Escherichia coli*, which belongs to the LysR-type transcriptional regulator (LTTR) family [15, 16]. This family of regulators is the most abundant in prokaryotes and features a conserved structure, with an N-terminal DNA-binding helix-turn-helix motif and a C-terminal co-inducer binding domain [16]. LTTRs are involved in sensing various chemical compounds, leading to the regulation of gene transcription through an allosteric-coupling mechanism central to biological systems [16]. HexA regulates several processes in *Photorhabdus*, including the production of secondary metabolites such as AQs and STs, bioluminescence, and cellular clumping [15–21]. However, the co-inducers for the LrhA family of regulators remain undefined [16].

*Photorhabdus* is the only Gram-negative bacterium known to biosynthesize stilbenes [15, 19]. Stilbene biosynthesis requires both cinnamic acid, resulting from the deamination of phenylalanine catalyzed by phenylalanine ammonia-lyase encoded by the *stlA* gene, and branched-chain fatty acids (BCFA) [22, 23]. The first stilbene derivative, 3,5-dihydroxy-4-isopropyl-trans-stilbene (IPS), results from the fusion of a cinnamic acid derivative and a BCFA precursor [22]. IPS, clinically known as tapinarof, is used as an anti-inflammatory agent in the treatment of psoriasis [3, 23–25]. IPS has also been shown to act as an antimicrobial, insect immunosuppressant, and signaling molecule in *P. laumondii*, inhibiting the production of AQ pigments and bioluminescence, and contributing to the mutualistic interaction within the nematode [22, 23, 26].

AQs are the largest group of natural pigments, most of which have been isolated from plants and are found in various human foods such as peas, cabbage, lettuce and beans [27]. Several biological activities have been reported for AQ-pigments, including antimalarial, antitumor, antioxidant, anti-inflammatory, and antimicrobial activities [27, 28]. The two *Photorhabdus* species, *P. laumondii* and *P. temperata,* are the only Gram-negative bacteria known to produce AQ-pigments via a type II polyketide-synthase (PKS) system, encoded by the biosynthesis gene cluster *antABCDEFGHI* [29, 30]. AntJ and CysB transcriptional regulators bind to the *antA* promoter, thereby activating AQ biosynthesis [30]. Eight different AQs have been described in various *Photorhabdus* strains [29, 31]. Most of these AQ molecules are derived from several *O-*methylations of the AQ-257 Da-derivative, involving homologous *O*-methyltransferase enzymes encoded in the genome of *P. laumondii* TT01 [31]. AQs are responsible for the yellow to red-orange pigmentation of bacterial colonies and cultures. However, their exact function in *Photorhabdus* life-cycle remains unclear [15, 30]. Nevertheless, some AQs exhibit weak antibiotic activity, leading to the suggestion that they may act as antagonistic agents against other microorganisms in the dead insect [7, 30, 32].

In the Gram-negative bacteria *E. coli*, *Salmonella,* and *P. laumondii*, the resistance-nodulation-division (RND)-type AcrAB-TolC efflux pump exports a wide range of substrates, including antibiotics, detergents, bile salts, dyes, bacterial metabolites, and other toxic compounds [33–37]. AcrAB also contributes to various physiological processes, such as detoxification, biofilm formation, virulence and motility, among others [37–40]. In our previous studies, we have shown that AcrAB is not essential for virulence but contributes to resistance to antimicrobial peptides (AMPs) such as polymyxins in *P. laumondii*. However, AcrAB exports the signaling molecule IPS, and its expression occurs during the stationary growth phase of *P. laumondii* [34, 35], concomitantly with the production and accumulation of secondary metabolites, particularly AQs and STs, in the insect cadaver [41].

In this study, we investigated the role of the AcrAB efflux pump in various phenotypic traits at both the transcriptional and targeted metabolite levels, focusing on AQs. RNA sequencing (RNA-seq) analysis revealed that the *acrAB* mutation alters the expression of genes involved in the biosynthesis of AQs, STs and fatty acids during the stationary growth phase. This mutation also negatively affected the expression of genes associated with motility during the exponential growth phase. Phenotypic analysis showed impaired motility, bioluminescence, and yellow-orange pigmentation in the Δ*acrA* mutant, whereas the Δ*stlA* mutant (deficient in stilbene production) exhibited increased bioluminescence and yellow-orange pigmentation. These results highlight the pleiotropic effects of the AcrAB efflux pump in *P. laumondii* TT01, potentially mediated in part through the export of signaling molecules such as IPS.

## Results and Discussion

### Deletion of *acrAB* strongly alters gene expression profile in *P. laumondii* TT01

In light of the known effects of IPS on *Photorhabdus* cellular processes [26] and its extrusion by AcrAB [34], we conducted a global transcriptomic study to explore the molecular impact of *acrAB* mutation *in P. laumondii*. The Δ*mdtA* mutant, which lacks the membrane fusion protein of the MdtABC RND-type efflux pump, was used as a negative control, as this pump does not exhibit any detrimental phenotypes in *Photorhabdus* [35, 42]. DESeq2 differential analysis comparing annotated features between the transcriptomes of different samples revealed no significant difference in gene expression between the WT and Δ*mdtA* mutant strains in either the exponential or stationary growth phases (Fig. S1A, Fig. S2 and Table S1A and B). Only 5 genes were significantly differentially expressed between the WT and Δ*mdtA* strains in the exponential growth phase, while 12 genes (1.6%) were significantly different in the stationary phase, including *mdtA*, *mdtB*, *mdtC* and *baeS* (Fig. S1A, Fig. S2 and Table S1A and B). In contrast, RNA-seq analysis revealed a significant difference in the expression of 43 genes, between the WT and the Δ*acrA* strains cultured to exponential growth phase (Fig. 1A, Fig. S1B and Table S2A). Twenty-five of these genes involved in fimbrial pilin and flagellar formation are downregulated (Fig. 1A, Fig. S1B and Table S2A). Notably, a significant difference in the expression of 185 genes (25.5%) was observed between the TT01 WT and Δ*acrA* grown to stationary growth phase (Fig. 1A and B, Fig. S1B, Fig. S2 and Table S2B). Among these, 115 genes were significantly downregulated, many of which are involved in the biosynthesis of STs (*stlC*, *stlD*, *stlE*, *stlA* and *plu2236* encoding a putative ST-epoxidase [13]), fatty acids (*plu2075*, *plu2217* and *plu2218*, encoding acyl-carrier proteins), AQs (*antABCDEFGHI*, *antJ* and *plu4892*-*plu4895,* encoding *O*-methyltransferase enzymes involved in *O*-methylations of the first of AQ derivative [31]), as well as genes related to bacteriophage tail fibers, cellular clumping, fimbrial pilin formation, and enterobactin biosynthesis (Fig. 1A and B, Fig. S2, and Table S2B).

**Fig. 1.**
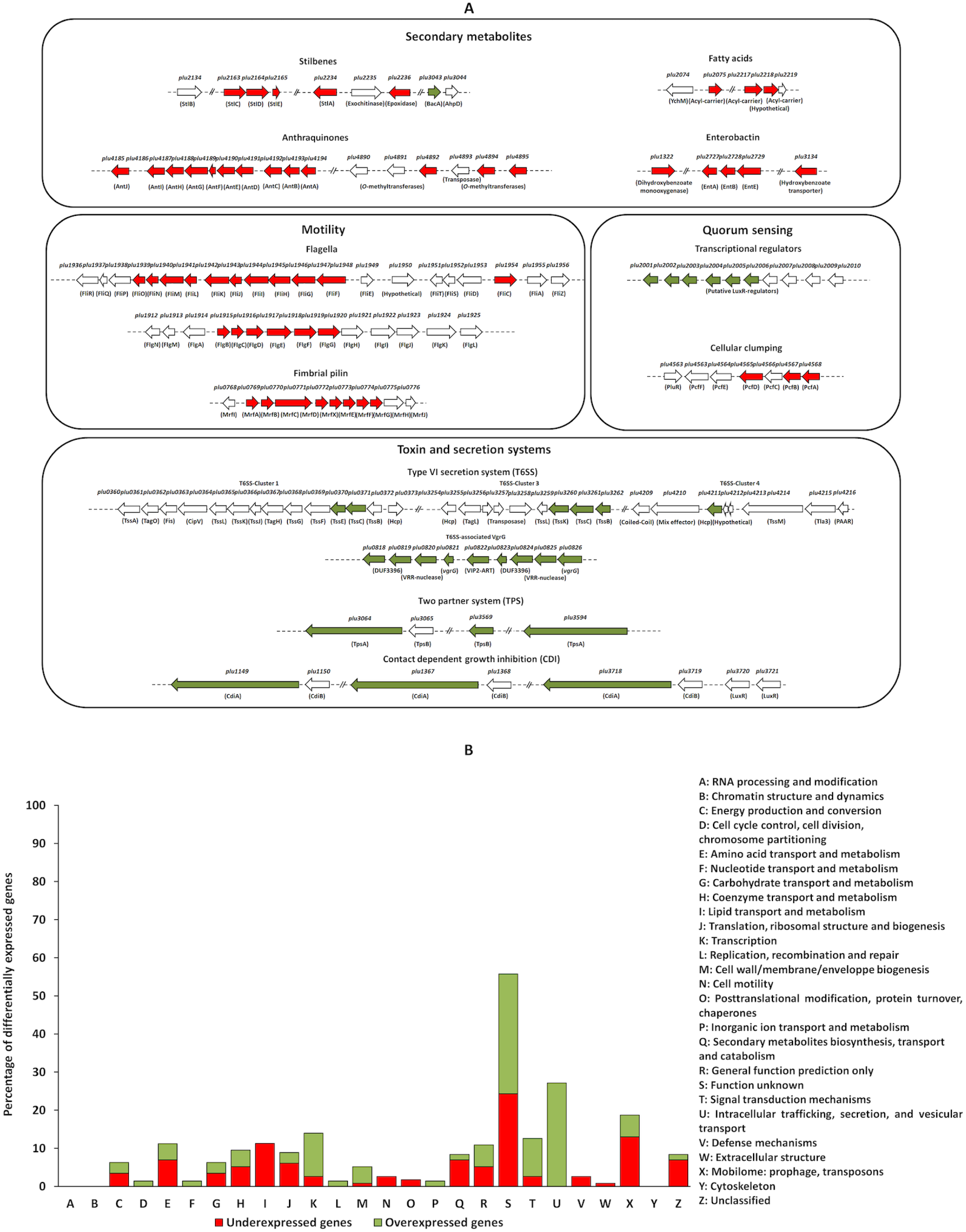
Identification of differentially expressed genes by RNA sequencing analysis between *P. laumondii* WT and Δ*acrA* mutant strains. *P. laumondii* WT and Δ*acrA* were cultured in LB medium to reach the exponential (OD_540_ = 0.5-0.7) and stationary (OD_540_ = 2.3-4.2) growth phases. (A) Functional classification of 82 of the 185 genes differentially expressed genes (|log_2_ fold change| ≥ 1, adjusted *p*-value (FDR) ≤ 0.05) into four groups: secondary metabolite biosynthesis, motility, quorum sensing, and toxin/secretion system biogenesis. Genes are represented by arrows labeled with their gene numbers and corresponding protein names. Downregulated genes are indicated by red arrows, upregulated genes by green arrows, and genes with similar expression levels between WT and Δ*acrA* are indicated with white arrows. (B) Clustering of 185 genes differentially expressed (|log_2_ fold change| ≥ 1, adjusted *p*-value (FDR) ≤ 0.05) between the WT and Δ*acrA* (EMBL accession number: BX470251 and GEO accession number: GSE280064), classified by Cluster of orthologous Groups (COG) annotation. The percentage of differentially expressed genes in each COG category is shown.

Conversely, 70 genes involved in T6SS (cluster 1, cluster 3 and cluster 4), T6SS-related VgrG, Tps-related proteins, CdiBA-related proteins, putative LuxR-type regulators, and other genes, such as those encoding Tcd insecticidal toxins, were upregulated in Δ*acrA* compared to WT (Fig. 1A and B, Fig. S2, and Table S2B). Interestingly, a Venn diagram analysis revealed that 61 common genes (8.6%) were differentially expressed in both a virulent polymyxin B resistant subpopulation of *P. laumondii* TT01 [43] and the Δ*acrA,* in comparison to the WT (Fig. S2). These genes include those encoding T6SS components, Tps-related proteins (*tpsA* and *tpsB*), CdiA effector proteins, and *pbgP4* encoding a PhoP-regulated protein involved in LPS modification and resistance to cationic AMPs. Both T6SS and CdiA toxins play crucial roles in bacterial competition, pathogenesis, and environmental stress survival [44–46]. Previous studies have shown that the polymyxin B resistant subpopulation is responsible for the virulence of *P. laumondii* TT01, that the *acrA* mutation did not significantly alter the percentage of the polymyxin B resistant subpopulation in *P. laumondii* TT01 [35]. These finding suggest that AcrAB is not involved in polymyxin B heteroresistance in *P. laumondii* TT01 and that the genes shared between Δ*acrA* and the polymyxin B-resistant subpopulation, which are differentially expressed involve other regulatory mechanisms.

Given that AcrAB plays a key role in the transport of various compounds across the bacterial membrane [37–40, 47], its involvement in extruding IPS signals [34, 35, 42], suggests that this efflux system may significantly impact the physiology of *Photorhabdus*. The exposure of *P. laumondii* WT to relevant concentrations of IPS induces a transcriptional stringent-like response in *Photorhabdus* [26], which could be further amplified in the AcrAB-mutant background, where intracellular IPS accumulates gradually. Indeed, our findings show that the differential gene expression profiles are specific to the *acrA* mutation in *P. laumondii* TT01, as no significant difference was detected between WT and Δ*mdtA* strain lacking the RND MdtABC efflux pump [42]. RT-qPCR confirmed this specificity showing a high correlation (0.98) with the RNA-seq data (Fig. S3A and B). These stress responses likely impair the biosynthesis of secondary metabolites, namely AQ and ST derivatives, by reallocating resources from secondary metabolism to stress survival mechanisms through feed-back loops.

Previous transcriptomic studies in other Enterobacterales such as *E. coli* and *Salmonella* showed that AcrAB primarily affects central metabolic pathways, such as carbohydrate metabolism, amino acid biosynthesis, the tricarboxylic acid (TCA) cycle, and fatty acid metabolism [37, 38]. This differs significantly from our findings in *Photorhabdus*, highlighting the versatility of AcrAB, which adapts its substrate profile in response to the physiological state and environmental conditions.

### AcrAB contributes to swarming and swimming motilities of *P. laumondii* TT01 in a stilbene independent manner

Several operons are associated with the structure and function of the flagella of *Photorhabdus*, thereby influencing the flagellum-driven motility of this bacterium [48]. These operons include the *motAB, flhBA, flg,* and *fli* loci in *P. laumondii* TT01 [49]. Our RNA-seq analysis revealed a downregulation of several genes related to flagellar function including *flgBCDEFG*, *fliFGHIJKLMNO,* and *fliC* in Δ*acrA* mutant strain grown to exponential growth phase compared to WT strain (Fig. 1A and Table S2A). However, no significant difference in expression levels of these genes was observed between WT and Δ*mdtA* strains (Table S1A and B). Using motility assays, we showed that Δ*acrA* exhibited reduced swarming and swimming motility compared to WT after 48 h and 24 h of incubation, respectively (Fig. 2A and B). No differences in the two types of motilities were observed between Δ*stlA* and the WT strain (Fig. 2A and B), demonstrating that motility in *P. laumondii* is IPS-independent. To test the potential involvement of RND-type efflux pumps in the export of other molecule(s), we tested the effect of phenyl-arginine-β-naphthylamide (PAβN), a competitive RND-type efflux pump inhibitor [42], on the swarming motility of *P. laumondii* strains. Our results demonstrated that PAβN strongly inhibited the swarming motility of all strains tested, including the Δ*acrA* mutant, after 48 h of incubation (Fig. 2A). Therefore, AcrAB likely contributes to *P. laumondii* motility by exporting signaling molecule(s) other than IPS, possibly by working with other efflux pumps. In contrast, our findings differ from a study on *S.* Typhimurium, which showed that motility-related genes, including *fli* and *flh* operons, were upregulated in an *acrB* mutant leading to enhanced motility. Recent studies have shown that endogenous metabolites such as polyamines produced by *E. coli* can bind to and activate AcrR, the transcriptional repressor of *acrAB*. Upon activation, AcrR is capable of binding to the promoter region of the *flhDC* operon encoding the master transcriptional regulator of the flagellar gene cascade, thereby repressing its expression [50, 51]. In *P. laumondii*, the reduced motility observed in Δ*acrA* and in all strains treated with PAβN may result from the impaired export of metabolites, which could negatively impact the bacterial fitness and, as a consequence, repress motility gene expression by binding accumulated metabolites to AcrR. An alternative explanation for the transcriptomic and phenotypic which affects key physiological functions in *S.* Typhimurium, such as motility [53]. Nevertheless, this hypothesis should also apply to all functional RND-type efflux pump including MdtABC in *P. laumondii* [35, 42]. However, our RNA-seq and phenotypic analyses showed no significant difference between Δ*mdtA* and TT01 WT, supporting that the observed transcriptional and phenotypic dysregulations in the Δ*acrA* would be indeed a consequence of a deficient AcrAB-dependent metabolite efflux.

**Fig. 2.**
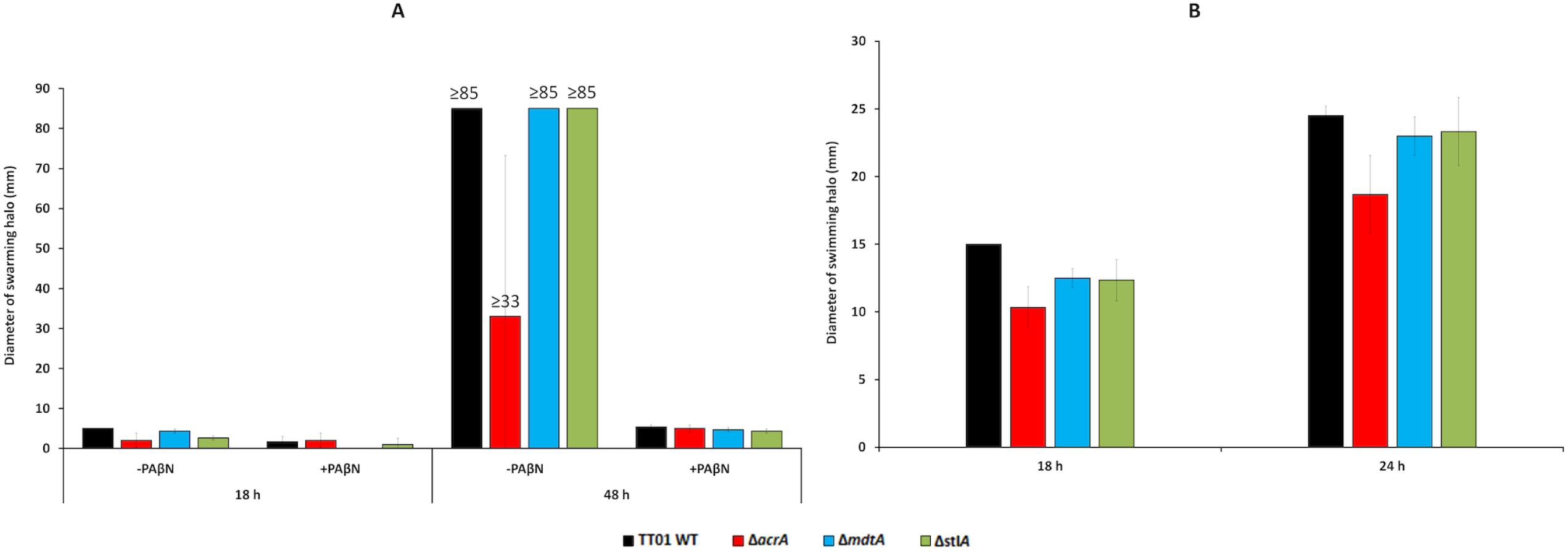
Swarming and swimming motilities of *P. laumondii* TT01 WT, Δ*acrA* and Δ*mdtA* strains. Diameters of motility halos were measured for three independent biological replicates of *P. laumondii* WT, Δ*acrA*, Δ*mdtA* and Δ*stlA* strains grown in LB broth supplemented with (A) 0.7% Eiken agar (swarming) with or without 50 mg.L^−1^ of phenyl-arginine-β-naphthylamide (PAβN) or (B) 0.35% agar (swimming). The cultures were incubated at 28°C for 48 h (swarming) or 24 h (swimming). Data are presented as mean ± standard errors of the mean (SEM).

### AcrAB contributes to bioluminescence production in a stilbene-dependent manner in *P. laumondii* TT01

Unlike many other Gram-negative bacteria, which communicate using acyl-homoserine-lactone (AHL) molecules, *Photorhabdus* employs pyrones for quorum sensing (QS). These pyrones are detected by the PluR regulator, a member of the LuxR-type regulators that control QS [20, 52, 53]. The PluR regulator directly binds to the promoter region of *pcfABCDEF* operon, encoding *Photorhabdus* Clumping Factor (Pcf) to activate cellular clumping in *P. laumondii* [20, 52]. Our RNA-seq analysis demonstrates a significant downregulation of *pcfAB* and *pcfD* genes involved in Pcf-mediated cellular clumping as well as up to 25 genes encoding tail fiber or bacteriophage related proteins in Δ*acrA* compared to WT when grown to stationary phase (Fig. 1A and B, and Table S2B). Clumping is thought to be primarily mediated by phage tail fibers, which likely function by forming bridges between bacterial cells [54]. Surprisingly, our cellular clumping assays revealed only a slight reduction in clumping of the Δ*stlA* strain relative to the WT, Δ*mdtA* and Δ*acrA* strains (Fig. S4). The global transcriptional regulator HexA has been identified as a repressor of cellular clumping in *P. laumondii,* by inhibiting the expression of the *pcf* operon [20]. Recently, it has been reported that different ligands, including IPS, which can act as a ligand for HexA [26], also compete for binding to the LuxR-type regulators involved in QS in several Gram-negative bacteria [55]. Taken together, these results suggest that HexA and LuxR regulators may compete for ligands such as IPS, which could mask a distinct phenotype of cellular clumping between TT01 WT and Δ*acrA*. This hypothesis is further supported by our RNA-seq analysis, which shows an upregulation of 6 genes encoding LuxR solo regulators, while *pluR* and 3 genes in the *pcf* operon—including *pcfC*, located between *pcfAB* and *pcfD*— were downregulated in the Δ*acrA* strain compared to WT (Fig. 1A and Table S2B).

QS, which is regulated by LuxR proteins, controls several processes, including bioluminescence in *Photorhabdus*—the only known example of a bioluminescent terrestrial bacterium [56, 57]. In *Photorhabdus*, bioluminescence is encoded by the *luxCDABE* operon, which governs luciferase-catalyzed oxidation of the luciferin substrate in presence of oxygen, resulting in the bioluminescent reaction [57]. The Δ*acrA* exhibited a reduction in bioluminescence intensity compared to the WT, Δ*mdtA* and Δ*stlA* strains, with a 3 h-delay compared to the other strains (Fig. 3). Interestingly, the bioluminescence intensity of Δ*stlA* was similar to all strains tested up to 14 h of incubation, when it becomes higher than that of WT, Δ*mdtA* and Δ*acrA*, particularly with a peak at 14 and 22 h of incubation (Fig. 3). No relevant difference in bioluminescence was observed between WT and Δ*mdtA* strains (Fig. 3). Surprisingly, our RNA-seq analysis revealed no transcriptional changes in the *lux* genes in the Δ*acrA* mutant compared to WT. Hapeshi and colleagues (2019) demonstrated that the exogenous addition of IPS to *P. laumondii* growing exponentially causes a rapid drop in bioluminescence, while the transcript levels of the *lux* operon itself were not affected [24]. Taken together, such data support the idea that bioluminescence in *Photorhabdus* is regulated at the post-transcriptional level. Indeed, the global transcriptional regulator HexA, inhibits bioluminescence at the post-transcriptional level [20, 56]. Since IPS, exported by AcrAB, may act as a cofactor of HexA and exogenous IPS rapidly reduces bioluminescence in *P. laumondii*, suggesting a role for AcrAB in modulating bioluminescence through HexA activation. Furthermore, the addition of PAβN to cultures led to a decrease in bioluminescence in all strains tested, with the Δ*acrA* mutant showing a more pronounced reduction compared to the WT, Δ*mdtA*, and Δ*stlA* strains, especially at 17 h of incubation (Fig. 3).

**Fig. 3.**
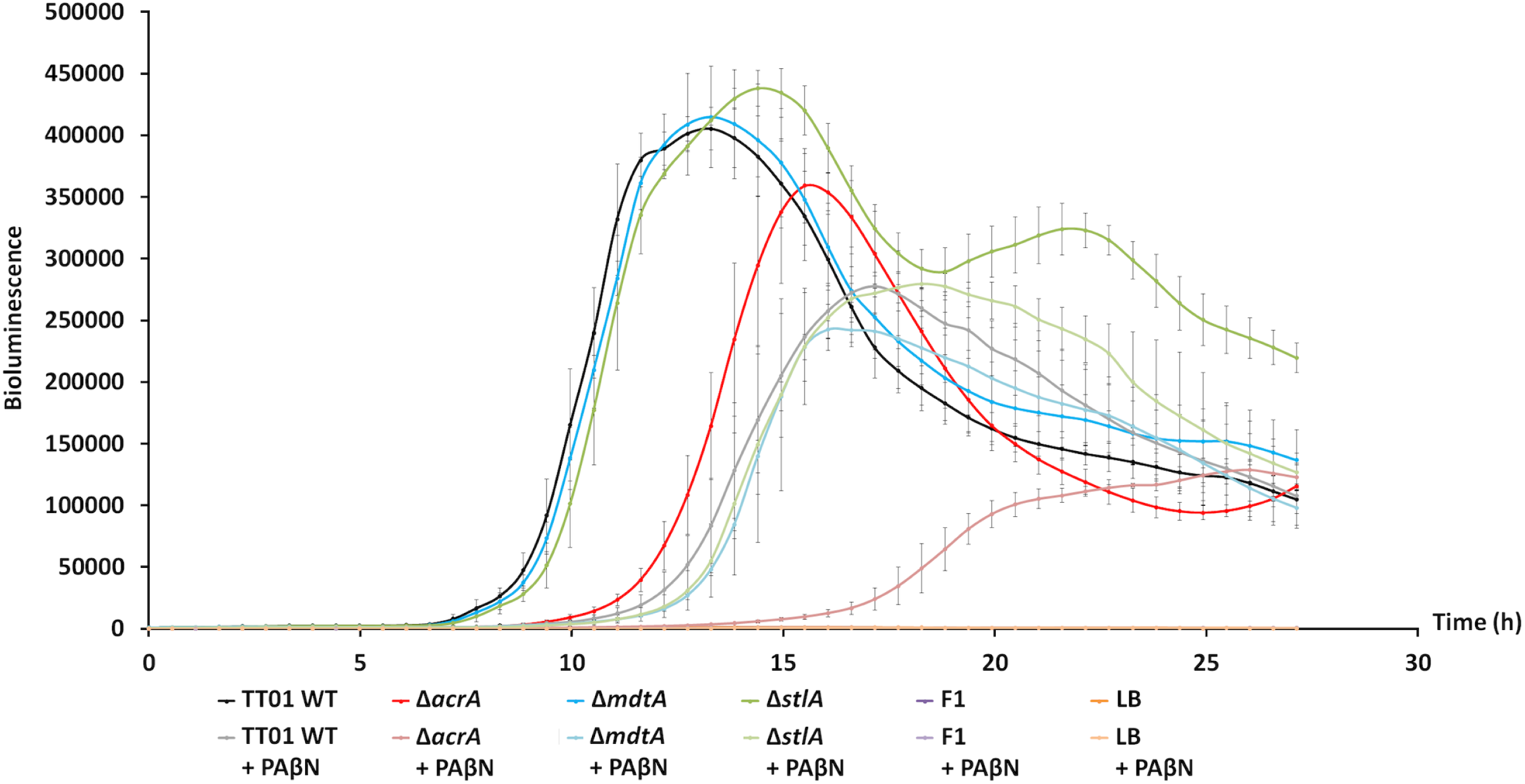
AcrAB contributes to the bioluminescence of *P. laumondii* TT01 in a stilbene-dependent manner. About 10^3^ of each *P. laumondii* WT, Δ*acrA*, Δ*mdtA*, and Δ*stlA* strains were inoculated into LB broth in 96-well white microtiter plates, with or without supplementation of 25 mg.L^−1^ PAβN, and grown for 27 h at 28°C. *Xenorhabdus nematophila* F1 and uninoculated LB broth were used as negative controls. Bioluminescence was monitored over time using an Infinite M200 microplate reader (Tecan). Data are from three independent biological and experimental replicates and are presented as means ± standard errors of the mean (SEM).

Taken together, these results suggest that AcrAB, exporting at least IPS as a signaling molecule, likely collaborates with other efflux pumps to modulate bioluminescence production in *P. laumondii* TT01.

### AcrAB contributes to the production of anthraquinone-mediated pigmentation in a stilbene-dependent manner in *P. laumondii* TT01

Our RNA-seq analysis revealed significant downregulation of 5 genes involved in the biosynthesis of STs, including *stlA*, *stlC*, *stlD*, and *stlE*, as well as 3 genes encoding acyl carrier proteins likely involved in ST production [22] (Fig. 1A). Additionally, 10 genes from the *antABCDEFGHI* operon, which is implicated in AQ biosynthesis, were downregulated, along with 3 genes (*plu4892*, *plu4894* and *plu4895*) encoding enzymes involved in AQ *O*-methylation (Fig. 1A, and Table S2B). Our phenotypic observations indicated that AQ production was impaired in the Δ*acrA* mutant, as evidenced by reduced pigmentation (white-beige colonies) compared to the typical yellow pigmentation of the WT strain on TSA (Fig. 4). The pigmentation defect in the Δ*acrA* mutant was restored upon introduction *in trans* of a wild-type copy of the *acrAB* operon (Fig. 4). In contrast, mutants lacking MdtABC or AcrAB-like RND-type efflux pumps [35, 42] did not exhibit pigmentation defects relative to the WT strain (Fig. 4). Interestingly, the Δ*stlA* mutant exhibited hyper-yellow to red-orange pigmentation, contrasting with the typical yellow pigmentation of the WT strain (Fig. 4). Consistent with this observation, a *P. laumondii* mutant strain defective in IPS production was previously shown to develop hyper-pigmented colonies on agar media [26]. Analysis of the absorbance spectra of extracts from cultures of WT, Δ*acrA*, Δ*acrA*/pBBR1-*acrA*, Δ*stlA* and Δ*stlA*/pBBR1-*stlA*, were performed using two commercial AQ molecules—Emodin (271 Da) and Physcion (285 Da)—as positive controls (absorbance peak 400-450 nm), revealed a decrease of AQ production by a factor of 2.5 in the Δ*acrA* mutant compared to both the WT and the Δ*acrA* mutant complemented with a WT *acrAB* operon. In contrast, the intensity of the absorbance at 400-450 nm was higher by a factor of 3.5 in the Δ*stlA* mutant compared to WT, suggesting increased AQ production in Δ*stlA* (Fig. 5A). Notably, the Δ*stlA* mutant strain complemented with a WT copy of the *stlA* gene displayed a decreased absorbance intensity at 400-450 nm by a factor of 6.8 compared to Δ*stlA* (Fig. 5A). These results are consistent with our RNA-seq analysis (Fig. 1A and Table S2B) and phenotypic observations on TSA plates (Fig. 4). Previous findings showed that *hexA* mutation activates IPS production but suppresses AQ production in *P. laumondii* [58], suggesting that in the absence of AcrAB efflux, IPS may activate HexA and inhibit the production of STs, while simultaneously inhibiting AQ production at the transcriptional level [26].

**Fig. 4.**
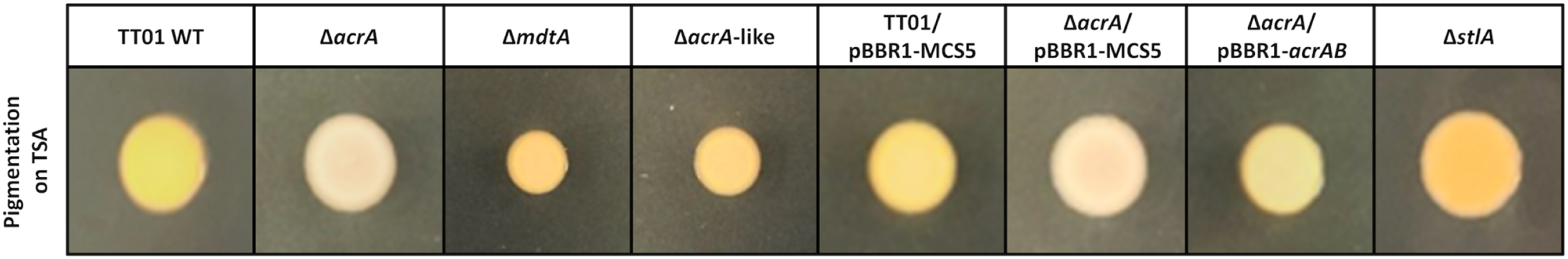
AcrAB contributes to anthraquinone-mediated pigmentation in *P. laumondii* TT01. Pigmentation was assessed on Trypticase Soja Agar (TSA) medium after 48 h of incubation at 28°C for each strain, harboring or not the pBBR1-MCS5 or the pBBR1-*acrAB* plasmids. Results shown are representative of at least three independent biological experiments, each with a minimum of three technical replicates.

**Fig. 5.**
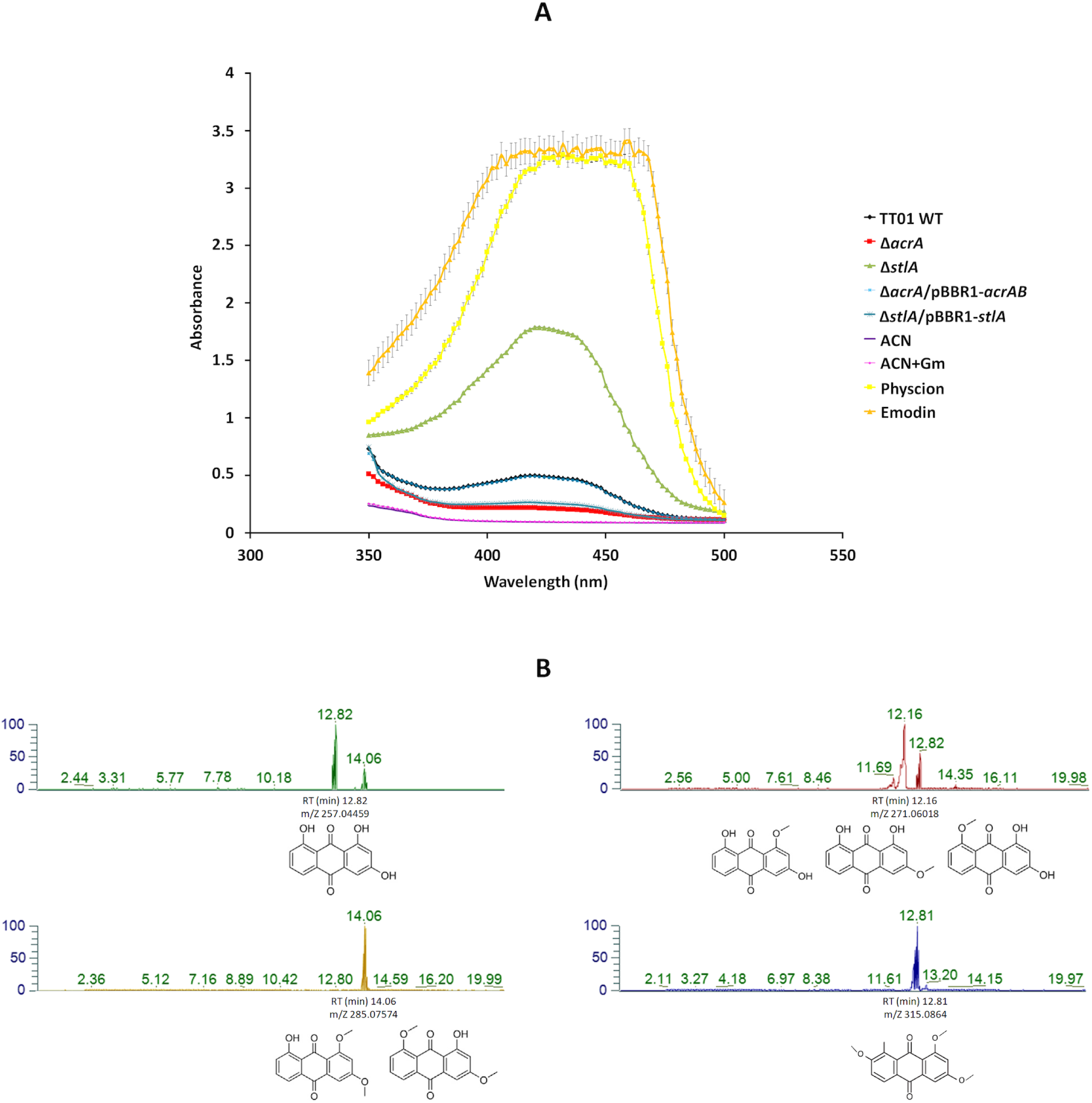
AcrAB is involved in anthraquinone-mediated yellow to red-orange pigmentation in *P. laumondii* TT01 strains in a stilbene-dependent manner. (A) Anthraquinone absorbance spectra were determined from culture extracts of *P. laumondii* WT, Δ*acrA*, Δ*acrA*/pBBR1-*acrAB*, Δ*stlA* and Δ*stlA*/pBBR1-*stlA* strains grown at 28°C in LB broth, supplemented with gentamicin at 15 mg.L^−1^ where appropriate. Absorbance measurements were recorded using an Infinite M200 microplate reader (Tecan). Acetonitrile (ACN) and acetonitrile + gentamycin (ACN + Gm) were used as negative control. Data represent averages ± standard errors of the mean (SEM) from three independent biological and experimental replicates. (B) Identification of anthraquinone derivatives was performed by extraction of targeted ion peak m/z chromatograms, analyzed by UHPLC-MS, from five independent extracellular and intracellular extracts of *P. laumondii* WT, Δ*acrA*, and Δ*stlA* strains grown in LB broth for 48 h at 28°C. A representative replicate of WT extracellular extracts is shown. Anthraquinone structures were adapted from [31] and modified from [29].

To confirm that the yellow to red-orange colors of *Photorhabdus* are due to AQ pigmentation, we screened several AQ derivatives produced by the WT strain. Previous reports have indicated that *Photorhabdus* primarily produces two AQ-derivatives, AQ 257 Da and AQ 271 Da, while AQ 301 Da and AQ 315 Da derivatives were exclusively identified in *Photorhabdus* cultured in insect larvae [31, 52]. In this study, UHPLC-MS analysis of extracts from 5 independent cultures of the WT, Δ*acrA* and Δ*stlA* strains in LB broth enabled us to identify 4 AQ derivatives naturally produced by the WT, with m/z [M+H]^+^ values of 257.045 Da, 271.060 Da, 285.075 Da and 315.086 Da, and retention times (t_R_) of approximatively 12.82, 12.16, 14.06 and 12.81 minutes, respectively (Fig. 5B and Fig. S5A, B, C and D). These were designated AQ 257 Da, AQ 271 Da, AQ 285 Da and AQ 315 Da (Fig. 5B and Fig. S5). While the localization of AQs in *P. laumondii* has not yet been determined, we performed two sequential intracellular extraction steps. The first step recovered intracellular molecules, while the second step specifically targeted molecules attached to the bacterial surface. The analysis of the extracts obtained from the intracellular and membrane fractions of WT, Δ*acrA*, and Δ*stlA* using UHPLC-MS showed that all detected AQ-derivatives were present in the supernatant, intracellular, and bacterial surface extracts (Fig. S5). These findings are in agreement with a study showing that AQ-derivatives may be present in different plant cell compartments [59], but have yet to be identified in *Photorhabdus*.

## Conclusion

Our study reveals an intricate interplay between the AcrAB-TolC efflux pump, specialized metabolite production, bioluminescence, and other physiological traits in *P. laumondii* TT01. AcrAB reduces the intracellular accumulation of toxic metabolites, including the small IPS molecule [34]. We demonstrate here that the transcriptional and phenotypic impacts of *acrAB* mutation on *P. laumondii* align with responses to ectopic IPS, involving genes regulated by HexA [20, 58], thereby confirming the role of IPS as signaling molecule. This influences profound gene expression changes and physiological responses in *Photorhabdus.* Therefore, we propose that *acrAB* mutation may cause an accumulation of intracellular IPS, which then acts as a ligand, enhancing the activity of the transcriptional inhibitor HexA and may cause export of as yet unidentified potential co-activator molecule(s), other than IPS, that could be involved in bacterial behaviors such as swarming and swimming motilities (Fig. 6 and Table 1).

**Fig. 6.**
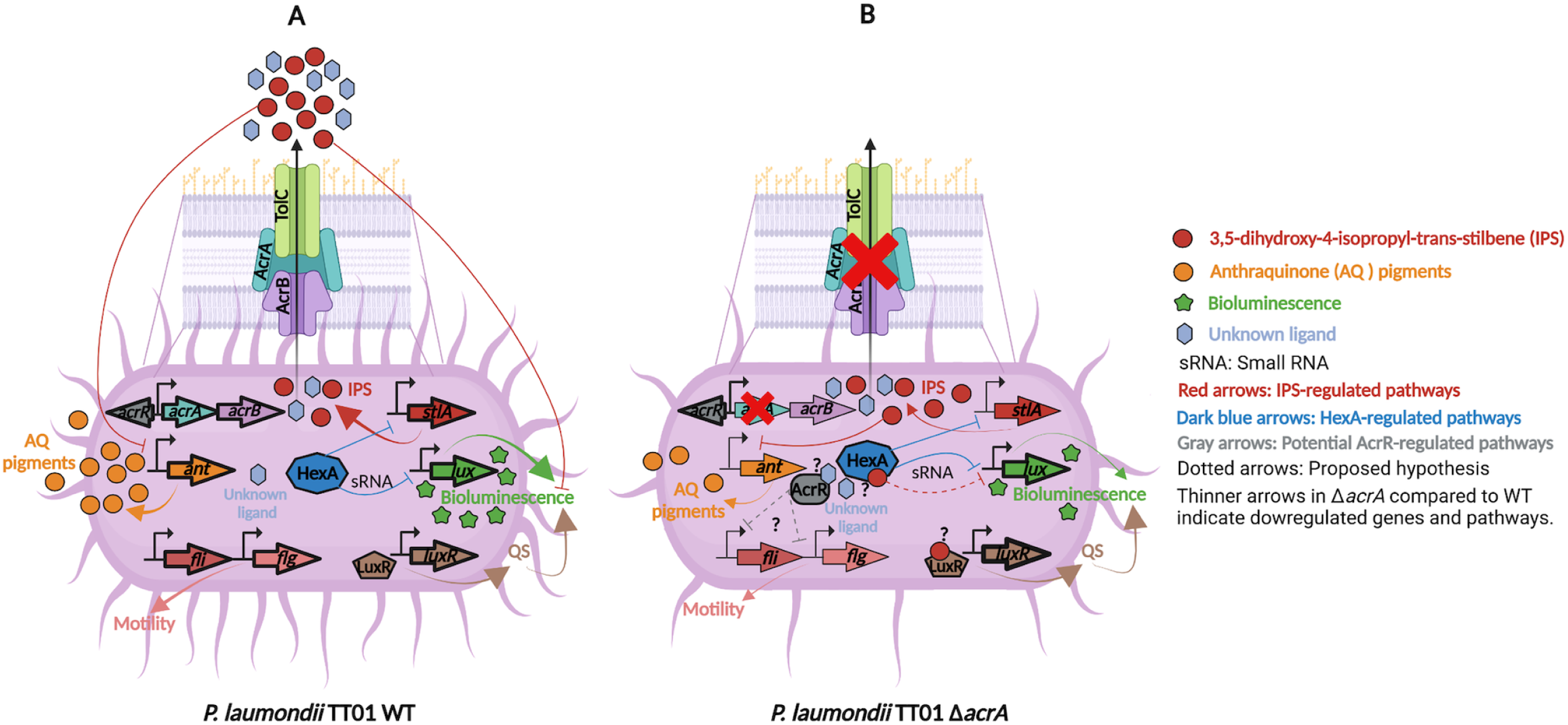
Proposed model for the pleiotropic effects of the AcrAB efflux pump in *P. laumondii* (A) WT and (B) Δ*acrA*. In *P. laumondii*, the *acrAB* operon encodes the AcrAB efflux pump, which consists of an inner membrane protein (AcrB), and a periplasmic fusion protein (AcrA). The *acrAB* operon is regulated by the local transcriptional repressor AcrR [33, 35]. The *stlA* gene is essential for stilbene biosynthesis [70], including IPS, which is exported by the AcrAB efflux pump in *P. laumondii* [34]. Furthermore, the exogenous addition of IPS inhibits (i) anthraquinone (AQ)-mediated yellow/orange pigmentation at the transcriptional level and (ii) bioluminescence in *P. laumondii* WT [26]. The *acrAB* mutation reduces the production of IPS, AQ, and bioluminescence. The global transcriptional regulator HexA exerts a negative feedback loop on stilbene biosynthesis, including IPS, and bioluminescence via small RNA [20, 58]. We propose that IPS may act as a ligand for HexA and LuxR-solos in *P. laumondii*. The potential accumulation of IPS in Δ*acrA* may (i) inhibit bioluminescence at the post-transcriptional level, as no difference in the expression level of the *luxCDABE* (*lux*) operon was detected compared to the WT, and (ii) upregulate LuxR-solos expression. AcrAB contributes to motility in *P. laumondii* WT independently of IPS. We showed that the *acrAB* mutation results in downregulation of several genes involved in motility in *P. laumondii*, leading to a reduced motility phenotype of Δ*acrA* compared to WT. We suggest that an unknown ligand, exported by AcrAB and potentially by other RND-type efflux pumps, may regulate the transcriptional levels of genes involved in motility, possibly via AcrR. This model was created using BioRender.com.

**Table 1.**
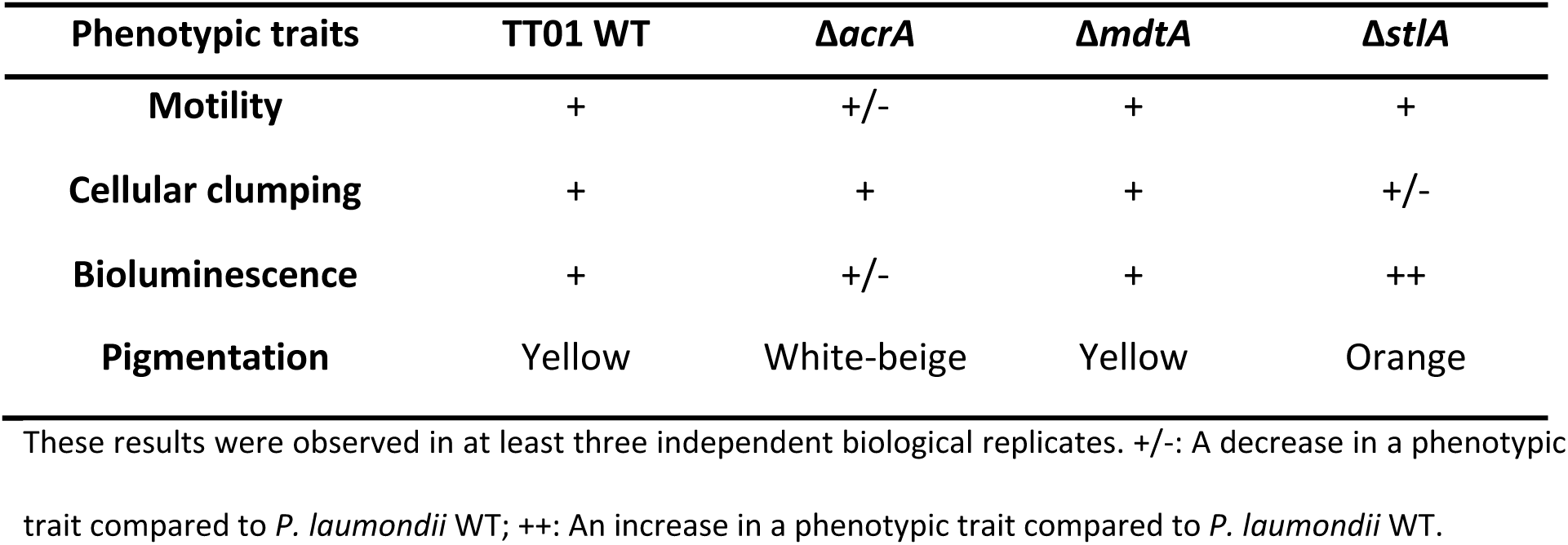
Phenotypic traits of different *P. laumondii* TT01 strains.

Overall, these findings highlight the multifaceted role of AcrAB in *P. laumondii*, extending beyond antibiotic resistance to include regulation of secondary metabolism and bacterial behavior. The regulatory connections between AcrAB, HexA, and the bacterial physiological responses underscore the complexity of bacterial regulatory networks. Future studies should elucidate the precise mechanisms by which AcrAB influences gene expression and to explore interactions between HexA, other transcriptional regulators and IPS. Additionally, the identification of other transcriptional regulator ligand(s) transported by RND-type efflux pumps will provide deeper insights into the regulatory architecture of Gram-negative bacteria.

## Materials and Methods

### Bacterial strains, plasmids and growth conditions

The bacterial strains and plasmids used in this study and their sources are listed in Table S3. *P. laumondii* TT01 [2], *Xenorhabdus nematophila* F1, *Escherichia coli* XL1-blue MRF’ (Stratagene) and WM3064 [60] were routinely grown in Luria Bertani (LB) broth or on agar (Difco) or in LB broth (Sigma) as previously described [34, 35]. When required, gentamicin (Gm) was added to LB broth/agar to a final concentration of 15 mg.L^−1^ for *E*. *coli* and *P. laumondii* TT01 strains harboring pBBR1-MCS5-derived plasmids.

### RNA extraction

Total RNA was extracted from bacterial strains grown in LB broth to exponential (OD_540_ = 0.5-0.7) and stationary (OD_540_ = 2.3-4.2) growth phases as previously described [42]. Nine independent biological replicates were performed per strain, and equal amounts of total RNA from three replicates of the same strain were pooled together. We thus generated three biological replicates per strain and condition that were subjected to rRNA depletion using the NEBNext® rRNA Depletion Kit (Bacteria) with RNA Sample Purification Beads prior to RNA sequencing.

### RNA sequencing (RNA-seq) and analysis

The eighteen RNA-seq libraries were performed and validated as described in [43]. Clustering and sequencing were performed on a NovaSeq 6000 (Illumina) on one lane of a flow cell SP.

Demultiplexing was performed with Illumina sequencing analysis software (CASAVA 1.8.2). Data quality was assessed with FastQC (Babraham Institute) and the sequencing analysis viewer (SAV) from Illumina software. Differential expression analysis was performed using the RNA-seq bioinformatics pipeline implemented in the MicroScope platform [61]. In a first step, forward raw RNA-seq reads of biological replicates samples were mapped onto the *P. laumondi* TT01 genome sequence (EMBL accession number: BX470251) as reference genome (BWA-MEM v.0.7.17) [62]. To take into account adapter sequences as well as poor quality sequences, an alignment score equal to at least half of the read length was required for a hit to be retained. The samtools suite (v.1.16.1) [63] was then used to extract reliable alignments with a Mapping Quality, MAPQ >= 1 from SAM files. From the 6 to 110 million Illumina sequences obtained, between 57% and 97% of high-quality mapped reads were finally kept. The number of these reads matching the genes of the reference genome was subsequently computed with bedtools coverageBed (v.2.30.0) [64]. Finally, the R package DESeq2 (v.1.22.2) [65] was used with lfcShrink function and its default parameters to normalize read counts and test for differential expression for each experimental conditions comparison. For each comparison, differential gene expression was considered significant for a computed adjusted *p*-value ≤ 0.05 and a log2 fold change ≥ 1 [66].

The complete data set from this study has been deposited in the NCBI’s Gene Expression Omnibus (GEO) database under accession number GSE280064.

### Quantitative RT-PCR (RT-qPCR) analysis

RT-qPCR was performed as previously described [34], using specific primer pairs for several selected genes (Table S4). For each reaction, melting curves were analyzed and each curve contained a single peak. Data were analyzed as the relative quantification ratio of a target gene versus the reference housekeeping gene *recA* between TT01 WT and *acrA* mutant strains using REST 2009 software as previously described [35]. *gyrB* gene was used as internal control.

### Swarming and swimming motility assays of *P. laumondii* TT01 strains

Swarming and swimming motility assays were determined by inoculating agar plates prepared with LB broth supplemented with 0.7% Eiken agar (Gerbu) or 0.35% bacto agar (Difco), respectively, with three independent biological replicates of bacterial strains, as previously described [67]. The diameters of the halos were measured 18 h, 24 h and 48 h after incubation at 28°C. The RND-type efflux pump inhibitor, phenyl-arginine-β-naphthylamide (PAβN, Sigma) was used at a final concentration of 50 mg.L^−1^.

Cellular clumping assays of *P. laumondii* TT01 strains.

Cell clumping assays were adapted from [20] and tested for TT01 WT, Δ*acrA*, Δ*mdtA* and Δ*stlA*. Three independent biological replicates of each culture were set to an optical density (OD_540_) of 0.05 and then incubated at 28°C without shaking in LB broth until exponential (OD_540_ = 0.28-0.4) and stationary (OD_540_ = 1.8-4.1) growth phases. Ten microliters of each culture were dropped onto 0.5% agarose pads (SeaKem GTG) in phosphate buffered saline (PBS) (Gibco) and then subjected to microscopic observation in contrast phase (Leica).

Bioluminescence measurements of *P. laumondii* TT01 strains.

Bioluminescence kinetics in LB broth were performed in white flat bottomed 96-well microtiter plates (Greiner) after inoculating about 10^3^ bacteria from an overnight culture of three independent biological replicates of bacterial strains into each microdilution well. *Xenorhabdus nematophila* F1 strain and LB broth medium were used as negative controls. Optical density (OD_540_) of all cultures was checked and bioluminescence was monitored for 27 h at 28°C with orbital shaking in an infinite M200 microplate reader (Tecan). Phenyl-arginine-β-naphthylamide (PAβN, Sigma) was used at a final concentration of 25 mg.L^−1^.

Pigmentation assays of *P. laumondii* TT01 strains.

Tinctorial pigmentation of different *P. laumondii* TT01 strains was revealed on Trypticase Soja Agar (TSA-bioMérieux). This medium is used for the detection of the yellow-orange AQs mediated pigmentation in *Photorhabdus* [1]. Five microliters of each bacterial cultures were spotted on TSA and then incubated at 28°C for 48 h in order for the colony pigmentation to be revealed.

### Determination of anthraquinone absorbance spectra in *P. laumondii* TT01 strains

Twenty-hour-old cultures from three independent replicates of bacterial strains grown in 5 mL LB broth (Sigma) supplemented with Gm when required, were used for inoculation with 1% (v/v) 10 mL of LB broth (Sigma). Cultures were incubated for 48 h at 28°C with shaking at 200 rpm. All cultures were checked for OD_540_, bacterial concentration and growth on nutrient bromothymol blue triphenyltetrazolium chloride agar (NBTA) medium as previously described [35]. Subsequently, three culture replicates from the same strain were pooled together and subjected to two successive ethyl acetate extractions (v/v) at room temperature for 24 h for each extraction to separate the organic phase. The extracts were evaporated, using a Stuart rotary evaporator connected to a water bath at 45°C (Fisher Scientific), followed by lyophilization. The resulting dry extracts were dissolved in 3 mL acetonitrile (ACN), and 200 µL/well of each extract was analyzed in three experimental replicates using an infinite M200 microplate reader (Tecan) in order to determine absorbance spectra at wavelengths between 350 and 500 nm. Two anthraquinone derivatives (Sigma): Emodin (271 Da) and Physcion (285 Da) dissolved in ACN at a final concentration of 500 mg.L^−1^ were used as positive controls. ACN with and without Gm were used as negative controls.

### Extracellular and intracellular extraction of anthraquinone pigments in *P. laumondii* TT01

Twenty-hour-old cultures from five independent replicates of bacterial strains cultured in 10 mL LB broth (Sigma), were used to inoculate at 1% (v/v) 100 mL of LB broth (Sigma). Cultures were incubated for 48 h at 28°C with shaking at 200 rpm. For all cultures, OD_540_, bacterial concentration and phenotypic traits (pigmentation aspect on TSA as described above and bacterial growth on NBTA medium as reported in [35]) were checked.

The extracellular extraction of AQ-pigments was performed after collecting supernatants by centrifugation each of 50 mL-cultures at 3500 x *g* for 30 min. Pellets were stored at −80°C to be used later for the intracellular-AQ extraction. We proceeded to two successive ethyl acetate extractions (v/v) at room temperature for 24 h. The extracts were evaporated, using a Stuart rotary evaporator connected to a water bath at 45°C (Fisher Scientific), followed by lyophilization.

Intracellular-AQ extraction protocols were adapted from [68, 69]. For each independent replicate, frozen bacterial pellet was suspended with 10 mL LB broth (Sigma) and sonicated for 6 min then cooled in ice every 1 min and 30 sec. Resulting suspensions were centrifuged for 20 min at 3500 x *g* in order to collect the supernatants (which correspond to the total intracellular content). Pellets were then suspended in 5 mL methanol (Sigma)/H_2_O 1:1 v/v with 1% formic acid (Sigma). All suspensions were subjected to ultrasonic treatment for 6 min and then centrifuged in order to collect supernatants. This final step allows for the collection of molecules that were attached to the bacterial surface. All supernatants resulted from intracellular extractions were lyophilized before being analyzed by UHPLC-MS.

### Characterization of anthraquinone pigments in *P. laumondii* TT01 by ultra-high-performance liquid chromatography-mass spectrometry (UHPLC-MS)

For UHPLC-MS analysis, we dissolved each extract with 2 mL of a methanol and Milli-Q water mix (50/50: v/v). After sample preparation, 10 μL of the supernatant of each replicate were injected onto a Thermo Vanquish Flex Binary UHPLC system coupled to a Q Exactive hybrid quadrupole-orbitrap mass spectrometer (Thermo Fisher Scientific, Waltham, MA, USA). The chromatographic separation was performed using an Aquity Premier BEH C18 column (1.7 μm, 2.1 × 100 mm, Waters, Milford, MA, USA) and two mobile phases: mobile phase A, 0.1% formic acid in Milli-Q water and mobile phase B, 0.1% formic acid in acetonitrile. The gradient was programmed as follows: 0–2 min, 100% A isocratic; 2–12 min, 0 to 50% B linear; 12–15 min, 50 to 100% B linear; 15–18 min, 100% B isocratic; 18–20 min, 100 to 0% B linear; followed by 2 min of re-equilibration of the column before the next run with a total run time of 20 min. The flow rate was set at 0.4 mL/min.

The UHPLC system was coupled to a Q Exactive™ Hybrid Quadrupole-Orbitrap High Resolution Mass Spectrometer (Thermo Fisher Scientific, San Jose, CA, USA). The MS acquisition was performed using positive ionization with a mass resolution of 120,000 at m/z 200. The m/z range for all full scan analyses was 200–1500. Heated electrospray ionization (HESI) parameters were as follows: sheath gas flow 60 arb (arbitrary units) auxiliary gas flow 15 arb, sweep gas flow 1 arb2 spray voltage 3.5 kV, probe temperature 400°C, capillary temperature 350°C. Prior to data collection, the mass spectrometer was calibrated using Pierce™ negative and positive ion calibration solution (Thermo Fisher Scientific, San Jose CA, USA). Row data were analyzed by FreeStyle™ 1.8 SP2 QF1 (Thermo Fisher Scientific, San Jose, CA, USA). Anthraquinones were characterized by their m/z [M+H]^+^ = 257.045 Da, 271.060 Da, 285.075 Da and 315.086 Da and t_R_ about 12.82, 12.16, 14.06 and 12.81 min, respectively [29–32].

## Acknowledgements

We thank, Philippe Clair from the BioCampus facility for his expert technical assistance with real-time PCR. MGX acknowledges the financial support provided by the France Génomique National infrastructure, funded under the “Investissement d’Avenir” program managed by the “Agence Nationale pour la Recherche” (contract ANR-10-INBS-09). Additionally, support from the “Plan France Relance” managed by the ANR (Molcheco Inrae/Nosopharm contract) is gratefully acknowledged. We also thank SPE, INRAE for awarding the IB2023 Scientific Grant. This work was also funded by a research grant from the Lebanese University (2022-2023) and a PHC CEDRE project 2022 (N47846 ZE).

